## Supplementary material for "Towards a unified molecular mechanism for ligand-dependent activation of NR4A-RXR heterodimers": Source Data: Nur77-RXRg Key Resources Table.docx

| **Key Resources Table** | | | | |
| --- | --- | --- | --- | --- |
| **Reagent type (species) or resource** | **Designation** | **Source or reference** | **Identifiers** | **Additional information** |
| strain, strain background (Escherichia coli) | BL21(DE3) | Sigma-Aldrich | CMC0014 | Electrocompetent cells |
| cell line (Homo-sapiens) | human embryonic kidney epithelial | ATCC | CRL-11268 |  |
| cell line (Homo-sapiens) | SK-N-BE(2)  neuroblastoma | ATCC | CRL-2271 |  |
| chemical compound, drug | BRF110 | This study |  | Synthesis procedure for BRF110 was described previously as cited in the methods. |
| chemical compound, drug | HX600 | Axon Medchem | CAS 172705- 89-4 |  |
| chemical compound, drug | 9-cis-Retinoic acid | Cayman Chemicals | CAS 5300-03-8 |  |
| chemical compound, drug | Bexarotene | Cayman Chemicals | CAS 153559- 49-0 |  |
| chemical compound, drug | LG100268 | Cayman Chemicals | CAS 153559-76-3 |  |
| chemical compound, drug | CD3254 | Cayman Chemicals | CAS 196961-43-0 |  |
| chemical compound, drug | SR11237 | Tocris Bioscience | CAS 146670-40-8 |  |
| chemical compound, drug | UVI3003 | Cayman Chemicals | CAS 847239-17-2 |  |
| chemical compound, drug | LG100754 | Cayman Chemicals | CAS 180713-37-5 |  |
| chemical compound, drug | IRX4204 | MedChemExpress | CAS 220619-73-8 |  |
| chemical compound, drug | Rhein | Sigma-Aldrich | CAS 478-43- 3 |  |
| chemical compound, drug | HX531 | Cayman Chemicals | CAS 188844-34-0 |  |
| chemical compound, drug | Danthron | Sigma-Aldrich | CAS 117-10-2 |  |
| chemical compound, drug | PA452 | Tocris Bioscience | CAS 457657-34-0 |  |
| peptide, recombinant protein | FITC-PGC1α | LifeTein |  | Amino acid sequence: EAEEPSLLKKLLLAPANTQ, with a N-terminal FITC label and an amidated C-terminus. |
| recombinant DNA reagent | Nur77-ligand binding domain (LBD) in pET45b(+) | This study | Bacteria expression plasmid | Residues: 356-598 |
| recombinant DNA reagent | RXRγ-ligand binding domain (LBD) in pET45b(+) | This study | Bacteria expression plasmid | Residues: 233-459 |
| recombinant DNA reagent | pET45b(+) | Novagen | 71327-3 |  |
| transfected construct (Photinus pyralis) | 3xNBRE-luciferase plasmid | de Vera et al., 2016 | Sanger sequenced |  |
| transfected construct (Photinus pyralis) | 3xDR1-luciferase plasmid | Hughes et al., 2014 | Mammalian expression plasmid, Sanger sequenced | This is the 3xPPRE-luciferase reporter plasmid in the referenced paper |
| transfected construct (human) | Full-length human Nur77 in pcDNA3.1 | This study | Mammalian expression plasmid, Sanger sequenced |  |
| transfected construct (human) | Full-length human RXRγ in pcDNA3.1 | This study | Mammalian expression plasmid, Sanger sequenced |  |
| recombinant DNA reagent | pcDNA3.1 empty vector | Thermo Fisher Scientific | V790-20 |  |
| sequence-based reagent | RXRγ-ΔLBD-F | This paper | PCR primer ordered from Sigma | CTACCAGTGGTTAGGAAGACATG |
| sequence-based reagent | RXRγ-ΔLBD-R | This paper | PCR primer ordered from Sigma | CATGTCTTCCTAACCACTGGTAG |
| sequence-based reagent | ΔNTD-RXRγ | This paper | gBlock sequences ordered from IDT | gBlock sequence in Supplementary file 1 |
| sequence-based reagent | RXRγ-hinge-LBD | This paper | gBlock sequences ordered from IDT | gBlock sequence in Supplementary file 1 |
| gene (human) | Nur77 (NR4A1) | Uniprot |  | Full length: residues 1-598; LBD: residues 356-598 |
| gene (human) | RXRγ (NR2B3) | Uniprot |  | Full length: residues 1-463; LBD: 226-462 |
| sequence-based reagent | Restriction enzymes, ligase for cloning | NEB |  | BamHI |
| sequence-based reagent | RXRγ-ΔNTD | This paper | gBlock for Gibson assembly | CTTAAGCTTGGTACCGAGCTCGATGTGTGCTATCTGTGGAGACAGATCCTCAGGAAAGCACTACGGGGTATACAGTTGTGAAGGCTGCAAAGGGTTCTTCAAGAGGACGATAAGGAAGGACCTCATCTACACGTGTCGGGATAATAAAGACTGCCTCATTGACAAGCGTCAGCGCAACCGCTGCCAGTACTGTCGCTATCAGAAGTGCCTTGTCATGGGCATGAAGAGGGAAGCTGTGCAAGAAGAAAGACAGAGGAGCCGAGAGCGAGCTGAGAGTGAGGCAGAATGTGCTACCAGTGGTCATGAAGACATGCCTGTGGAGAGGATTCTAGAAGCTGAACTTGCTGTTGAACCAAAGACAGAATCCTATGGTGACATGAATATGGAGAACTCGACAAATGACCCTGTTACCAACATATGTCATGCTGCTGACAAGCAGCTTTTCACCCTCGTTGAATGGGCCAAGCGTATTCCCCACTTCTCTGACCTCACCTTGGAGGACCAGGTCATTTTGCTTCGGGCAGGGTGGAATGAATTGCTGATTGCCTCTTTCTCCCACCGCTCAGTTTCCGTGCAGGATGGCATCCTTCTGGCCACGGGTTTACATGTCCACCGGAGCAGTGCCCACAGTGCTGGGGTCGGCTCCATCTTTGACAGAGTCCTAACTGAGCTGGTTTCCAAAATGAAAGACATGCAGATGGACAAGTCGGAACTGGGATGCCTGCGAGCCATTGTACTCTTTAACCCAGATGCCAAGGGCCTGTCCAACCCCTCTGAGGTGGAGACTCTGCGAGAGAAGGTTTATGCCACCCTTGAGGCCTACACCAAGCAGAAGTATCCGGAACAGCCAGGCAGGTTTGCCAAGCTGCTGCTGCGCCTCCCAGCTCTGCGTTCCATTGGCTTGAAATGCCTGGAGCACCTCTTCTTCTTCAAGCTCATCGGGGACACCCCCATTGACACCTTCCTCATGGAGATGTTGGAGACCCCGCTGCAGATCACCTGAGATCCACTAGTCCAGTGTGG |
| sequence-based reagent | RXRγ-hinge-LBD | This paper | gBlock for Gibson assembly | CTTAAGCTTGGTACCGAGCTCGATGAAGAGGGAAGCTGTGCAAGAAGAAAGACAGAGGAGCCGAGAGCGAGCTGAGAGTGAGGCAGAATGTGCTACCAGTGGTCATGAAGACATGCCTGTGGAGAGGATTCTAGAAGCTGAACTTGCTGTTGAACCAAAGACAGAATCCTATGGTGACATGAATATGGAGAACTCGACAAATGACCCTGTTACCAACATATGTCATGCTGCTGACAAGCAGCTTTTCACCCTCGTTGAATGGGCCAAGCGTATTCCCCACTTCTCTGACCTCACCTTGGAGGACCAGGTCATTTTGCTTCGGGCAGGGTGGAATGAATTGCTGATTGCCTCTTTCTCCCACCGCTCAGTTTCCGTGCAGGATGGCATCCTTCTGGCCACGGGTTTACATGTCCACCGGAGCAGTGCCCACAGTGCTGGGGTCGGCTCCATCTTTGACAGAGTCCTAACTGAGCTGGTTTCCAAAATGAAAGACATGCAGATGGACAAGTCGGAACTGGGATGCCTGCGAGCCATTGTACTCTTTAACCCAGATGCCAAGGGCCTGTCCAACCCCTCTGAGGTGGAGACTCTGCGAGAGAAGGTTTATGCCACCCTTGAGGCCTACACCAAGCAGAAGTATCCGGAACAGCCAGGCAGGTTTGCCAAGCTGCTGCTGCGCCTCCCAGCTCTGCGTTCCATTGGCTTGAAATGCCTGGAGCACCTCTTCTTCTTCAAGCTCATCGGGGACACCCCCATTGACACCTTCCTCATGGAGATGTTGGAGACCCCGCTGCAGATCACCTGAGATCCACTAGTCCAGTGTGG |
| commercial assay or kit | Gibson assembly | NBE | E2611L |  |
| commercial assay or kit | Britelite plus Reporter Gene Assay System | Perkin Elmer | 6066769 |  |
| commercial assay or kit | LanthaScreen Elite Tb-anti-His antibody | Thermo Fisher | #PV5895 |  |
| software, algorithm | NITPIC software | Keller et al., 2012 |  | Baseline calculation, curve integration |
| software, algorithm | SEDPHAT | Brautigam et al., 2016 |  | Estimation of binding affinity and thermodynamic parameter measurements |
| software, algorithm | GUSSI | Brautigam et al., 2015 |  | Plot ITC figures |
| other | NMR chemical shift assignment of Nur77 LBD | This paper | BMRB52973 | Published NMR peak assignment from Biological Magnetic Resonance Data Bank |
| software, algorithm | NMRFx | Norris et al., 2016 |  | NMR data process and analysis |
| software, algorithm | Pearsons and Spearman correlation analysis | GraphPad Prism |  | Correlation analysis |
| software, algorithm | Principal Component Analysis (PCA); | GraphPad Prism |  | Correlation analysis |
| software, algorithm | ANOVA multiple comparison test | GraphPad Prism |  | Statistical testing |
